# Vitamin D status and risk of incident tuberculosis disease: a systematic review and individual participant data meta-analysis

**DOI:** 10.1101/481739

**Authors:** Omowunmi Aibana, Chuan-Chin Huang, Said Aboud, Alberto Arnedo-Pena, Mercedes C. Becerra, Juan Bautista Bellido-Blasco, Ramesh Bhosale, Roger Calderon, Silvia Chiang, Carmen Contreras, Ganmaa Davaasambuu, Wafaie W. Fawzi, Molly F. Franke, Jerome T. Galea, Daniel Garcia-Ferrer, Maria Gil-Fortuño, Barbará Gomila-Sard, Amita Gupta, Nikhil Gupte, Rabia Hussain, Jesus Iborra-Millet, Najeeha T. Iqbal, Jose Vicente Juan-Cerdán, Aarti Kinikar, Leonid Lecca, Vidya Mave, Noemi Meseguer-Ferrer, Grace Montepiedra, Ferdinand M. Mugusi, Olumuyiwa A. Owolabi, Julie Parsonnet, Freddy Roach-Poblete, Maria Angeles Romeu-García, Stephen A. Spector, Christopher R. Sudfeld, Mark W. Tenforde, Toyin O. Togun, Rosa Yataco, Zibiao Zhang, Megan B. Murray

**Affiliations:** Department of Internal Medicine, McGovern Medical School at the University of Texas Health Science Center, 6431 Fannin Street, MSB 1.122, Houston, Texas 77030, USA (O Aibana MD); Department of Global Health and Social Medicine, Harvard Medical School, 641 Huntington Avenue, Boston, Massachusetts 02115, USA (C-C Huang ScD); Department of Microbiology and Immunology, Muhimbili University of Health and Allied Sciences, United Nations Road, Upanga West, Dar es Salaam, Tanzania (S Aboud MD); Epidemiology Division, Public Health Center, C/L’Olivera 5, 2-C. 12005, Castellon, Spain (A Arnedo-Pena MD); Department of Global Health and Social Medicine, Harvard Medical School Boston, 641 Huntington Avenue, Massachusetts 02115, USA (M C Becerra ScD); Epidemiology Division, Public Health Center, Avenida del Mar 8, 12003, Castellon, Spain (J Bellido-Blasco MD); Department of Obstetrics & Gynecology, Byramjee Jeejeebhoy Government Medical College, Pune, 411001, India (R Bhosale MD); Partners in Health - Socios En Salud Sucursal, Av. Chimpu Ocllo # 998 Urb. Residencial Lucyana, Carabayllo, Lima, Peru (R Calderon MPH); Department of Pediatrics, Alpert Medical School of Brown University, 55 Claverick Street, Suite 101, Providence, Rhode Island, USA (S Chiang MD); Partners in Health - Socios En Salud Sucursal, Calle Los Geranios 312, Lince, Lima, Peru (C Contreras MS); Channing Division of Network Medicine, Department of Medicine, Brigham and Women’s Hospital, Harvard Medical School, 665 Huntington Avenue, Boston, Massachusetts 02115, USA (G Davaasambuu PhD); Department of Global Health and Population, Harvard T.H. Chan School of Public Health, 665 Huntington Avenue, Boston, Massachusetts 02115, USA (W W Fawzi DrPH); Department of Global Health and Social Medicine, Harvard Medical School, 641 Huntington Avenue, Boston, Massachusetts 02115, USA (M F Franke ScD); School of Social Work, University of South Florida, 13301 Bruce B Downs Blvd, MHC 1416 A, Tampa, Florida 33612, USA (J T Galea PhD); Biochemical Laboratory, Hospital General Castellon, Spain, C/R.O. Caballeros del Puig 5, 8. 46540 El Puig, Valencia, Spain (D Garcia-Ferrer PharmD); Microbiology Laboratory, Hospital General Castellon, Spain, C/Monastir de Poblet 17, 5K. 12540 Vila-real, Castellon, Spain (M Gil-Fortuño MD); Microbiology Laboratory, Hospital General Castellon, Spain, Residencial Montemar 8. 46112 Massarrochos, Valencia Spain (B Gomila-Sard MD); Division of Infectious Diseases, Department of Medicine, Johns Hopkins University School of Medicine, 600 North Wolfe Street, Baltimore, Maryland 21287, USA (A Gupta MD); Byramjee Jeejeebhoy Government Medical College-Johns Hopkins University CRS, Jai Prakash Narayan Road, Pune 411001, India (N Gupte PhD); Department of Pathology and Microbiology, Aga Khan University, Stadium Road, P.O. BOX 3500, Karachi, Pakistan (R Hussain PhD); Biochemical Laboratory, Hospital General Castellon, Spain, C/ Puig 12, BJ. 46138 Rafelbuñol (Valencia) Spain (J Iborra-Millet; PharmD); Department of Pediatrics and Child Health & Biological and Biomedical Sciences, Stadium Road P.O. BOX 3500, Karachi, Pakistan (N T Iqbal PhD); Biochemical Laboratory, Hospital General Castellon, Spain, Avenida Burjasot 100, 9. 46009 Valencia Spain (J Juan-Cerdán MD); Department of Pediatrics. Byramjee Jeejeebhoy Government Medical College, Jai Prakash Narayan Road, Near Pune Railway Station, Pune 411001, India (A Kinikar MD); Partners in Health - Socios En Salud Sucursal, Los Geranios 312, Lima, Peru (L Lecca MD); Byramjee Jeejeebhoy Government Medical College-Johns Hopkins University CRS, Jai Prakash Narayan Road, Pune 411001, India (V Mave MD); Epidemiology Division, Public Health Center, C/San Roque 84, 6-11. 12004, Castellon, Spain (N Meseguer-Ferrer, RN); Center for Biostatistics in AIDS Research & Department of Biostatistics, Harvard T.H. Chan School of Public Health, Bagnoud Bldg, CBAR FXB-5th Flr, 651 Huntington Ave, Boston, Massachusettes USA 02115 (G Montepiedra PhD); Department of Internal Medicine, Muhimbili University of Health and Allied Sciences, United Nations Road, Upanga West, Dar es Salaam, Tanzania (F M Mugusi MD); Medical Research Council Gambia at London School of Hygiene and Tropical Medicine, PO Box 273, Banjul, The Gambia (O A Owolabi MD); Departments of Medicine and of Health Research and Policy, Stanford University School of Medicine, Lane L134, 300 Pasteur Drive, Stanford, California 94305, USA (J Parsonnet MD); Laboratory Hospital Regional Antofagasta, General Pedro Lagos 253. Playa Blanca, Antofagasta, Chile (F Roach-Poblete MD); Epidemiology Division, Public Health Center, C/Bayer 6, 2. 12002, Castellon, Spain (M Romeu-García MD); Division of Infectious Diseases, Department of Pediatrics, University of California San Diego, 9500 Gilman Drive, La Jolla, California 92093, USA (S A Spector MD); Department of Global Health and Population, Harvard T.H. Chan School of Public Health, 665 Huntington Avenue, Boston, Massachusetts 02115, USA (C R Sudfeld ScD); Division of Allergy and Infectious Diseases, Department of Medicine, University of Washington School of Medicine, 1959 NE Pacific Street, Seattle, Washington 98195, USA (M W Tenforde MD); Department of Epidemiology and Biostatistics, McGill University, 1020 Pine Avenue West, Montreal, Quebec H3A 1A2, Canada (T O Togun PhD); Partners in Health - Socios En Salud Sucursal, Av. San Luis 1040 Condominio del Aire Block S2, Dpt 502, San Luis, Lima, Peru (R Yataco BS); Division of Global Health Equity, Brigham and Women’s Hospital, Harvard Medical School, 641 Huntington Avenue, Boston, Massachusetts 02115, USA (Z Zhang MS); Department of Global Health and Social Medicine, Harvard Medical School, 641 Huntington Avenue, Boston, MA 02115, USA (Professor M B Murray MD)

**Author notes:** **Correspondence to**: Dr Megan B. Murray, Department of Global Health and Social Medicine, Harvard Medical School, 641 Huntington Avenue, 4th Floor, Room 4A07, Boston, MA 02115.

## Abstract

**Background:** Few studies have evaluated the association between pre-existing vitamin D deficiency (VDD) and incident TB. We assessed the impact of baseline vitamin D on TB risk.

**Methods:** We assessed the association between baseline vitamin D and incident TB in a prospective cohort of 6751 household contacts of TB patients in Peru. We also conducted a one-stage individual participant data meta-analysis searching PubMed and Embase for studies of vitamin D and TB until December 31, 2017. We included studies that assessed vitamin D before TB diagnosis. We defined VDD as 25–(OH)D <50 nmol/L, insufficiency as 50–75 nmol/L and sufficiency as >75nmol/L. We estimated the association between vitamin D and incident TB using conditional logistic regression in the Peru cohort and generalized linear mixed models in the meta-analysis.

**Findings:** In Peru, baseline VDD was associated with a statistically insignificant increase in incident TB (aOR 1·70, 95% CI 0·84–3·46; p=0·14). We identified seven studies for the meta-analysis and analyzed 3544 participants. Individuals with VDD and very low vitamin D (<25nmol/L) had increased TB risk (aOR 1·48, 95% CI 1·04–210;*p*=0· 03 and aOR 2 08, 95% CI 0·88–4·92; *p* trend=002 respectively). Among HIV-positive patients, VDD and very low vitamin D conferred a 2-fold (aOR 2.18, 95% CI 1· 22–3·90; p=0· 01) and 4-fold (aOR 4·28, 95% CI 0·85–21·44; p trend=0·01) increased risk of TB respectively.

**Interpretation:** Our findings suggest vitamin D predicts TB risk in a dose-dependent manner and vitamin D supplementation may play a role in TB prevention.

**Funding:** National Institute of Health (NIH), National Institute of Allergy and Infectious Diseases (NIAID), National Institute on Drug Abuse (NIDA), National Institute of Mental Health (NIMH), International Maternal Pediatric Adolescent AIDS Clinical Trials Group (IMPAACT), Eunice Kennedy Shriver National Institute of Child Health and Human Development (NICHD), National Institute of Dental and Craniofacial Research (NIDCR), Boehringer-Ingelheim, Bristol-Myers Squibb, Gilead Sciences, GlaxoSmithKline, Foundation, Ujala Foundation, Wyncote Foundation, NIH - Fogarty International Center Program of International Training Grants in Epidemiology Related to AIDS, NIAID Byramjee Jeejeebhoy Medical College HIV Clinical Trials Unit, NIAID’s Baltimore-Washington-India Clinical Trials Unit, National Commission on Biotechnology, the Higher Education Commission, International Research Support Initiative Program of the Higher Education Commission Government of Pakistan, the Bill and Melinda Gates Foundation, and the NIH Fogarty International Center.

**Research in Context:** *Evidence before this study:* Numerous studies have found lower serum vitamin D levels among patients with active TB disease compared to healthy controls. However, research has not clarified whether low vitamin D increases TB risk or whether TB disease leads to decreased vitamin D levels. We conducted PubMed and Medline searches for all studies available through December 31, 2017 on the association between vitamin D status and TB disease. We included the following keywords: “vitamin D,” “vitamin D deficiency,” “hypovitaminosis D,” “25-hydroxyvitamin D,” “1,25-dihydroxyvitamin D,” “vitamin D2,” “vitamin D3,” “ergocalciferol,” “cholecalciferol,” and “tuberculosis.” We found only seven studies had prospectively evaluated the impact of baseline vitamin D levels on risk of progression to TB disease. We report here the results of a case control study nested within a large prospective longitudinal cohort study of household contacts of TB cases and the results of an individual participant data (IPD) metaanalysis of available evidence on the association between vitamin D levels and incident TB disease.

*Added value of this study:* We demonstrated that low vitamin D levels predicts risk of future progression to TB disease in a dose-dependent manner.

*Implications of all the available evidence:* These findings suggest the possibility that vitamin D supplementation among individuals at high risk for developing TB disease might play a role in TB prevention efforts.

## Introduction

The global burden of tuberculosis (TB) remains high with approximately one-fourth to one-third of world’s population infected with *Mycobacterium tuberculosis*, and the World Health Organization (WHO) estimates 10 million people developed TB disease in 2017.^1^ Concurrently, vitamin D deficiency (VDD) is a widespread problem, with reported adult prevalence of VDD ranging from 10% to 80% worldwide.^2,3^ Vitamin D is an important regulator of the immune system,^4^ and *in vitro* studies have elucidated some of the mechanisms by which vitamin D influences TB disease pathogenesis.^5,6^ The discovery that vitamin D activates cathelicidin-mediated killing of ingested mycobacteria in macrophages^5^ has focused attention on the possibility that low vitamin D levels may contribute to TB disease progression.

Numerous observational studies have also documented lower serum vitamin D levels among TB patients compared to healthy controls, and prior meta-analyses investigating the association between vitamin D and TB have concluded that low vitamin D increases TB disease risk.^7–10^ However, most studies were cross-sectional studies and assessed vitamin D status after the diagnosis of active TB disease, rather than the impact of pre-existing vitamin D levels on the risk of progression to TB disease. Given TB disease can induce profound metabolic abnormalities, it remains unclear whether vitamin D deficiency increases TB disease risk or whether TB disease leads to decreased serum vitamin D levels. Furthermore, prior studies evaluating the association between vitamin D and TB disease have used different cutoffs to categorize vitamin D levels or define vitamin D deficiency.^7–10^ Hence, it is challenging to determine whether there is a vitamin D threshold below which individuals are at risk of TB disease.

Here, we address the association of vitamin D levels on the risk of TB progression in two ways. We first report results of a case control analysis nested in a prospective cohort study of household contacts of TB patients that we conducted in Lima, Peru. We next pool these data with those from other published prospective studies of vitamin D status and TB risk to conduct an individual participant data (IPD) meta-analysis synthesizing available evidence on the association between vitamin D levels and incident TB disease.

## Methods

### Lima Cohort Study

#### Ethics Statement

The study was approved by the Institutional Review Board of Harvard School of Public Health and the Research Ethics Committee of the National Institute of Health of Peru. All study participants or guardians provided written informed consent.

#### Study Setting and Population

Details of the study design and methods are described elsewhere^11,12^ and presented in the appendix (p 1). We enrolled a prospective cohort of household contacts (HHCs) of index TB patients in Lima, Peru between September 2009 and August 2012. We screened HHCs for pulmonary and extra-pulmonary TB disease at baseline and 2, 6, and 12 months after enrollment. We offered HIV testing and invited HHCs to provide a baseline blood sample. We classified HHCs as having incident secondary TB disease if they were diagnosed at least 15 days after index case enrollment and co-prevalent TB disease if they were diagnosed earlier.

We defined “cases” as HIV-negative HHCs with blood samples who developed incident secondary TB disease within one year of follow-up. For each case, we randomly selected four controls from among HHCs who were not diagnosed with TB disease, matching on gender and age by year.

At the end of follow up, we measured levels of total 25–(OH)D and retinol from stored blood samples.

#### Statistical Analysis

We defined vitamin D deficiency as serum 25–(OH)D < 50 nmol/L, insufficiency as 50–75 nmol/L and sufficiency as > 75nmol/L.^13^ We used univariate and multivariate conditional logistic regression models to evaluate the association between baseline VDD and risk of TB disease. Because we had previously observed that vitamin A deficiency increases TB risk in this cohort,^11^ we also adjusted for vitamin A levels.

In sensitivity analyses, we restricted the analysis to HHCs diagnosed with incident TB at least 60 days after index patient enrollment and their matched controls. We also separately evaluated patients with microbiologically confirmed TB and their matched controls.

### Systematic Review and IPD Meta-analysis

#### Search Strategy and Data Sources

We conducted the systematic review and meta-analysis according to the Preferred Reporting Items for Systematic Reviews and Meta-Analyses (PRIMSA) guidelines.^14,15^ All studies included in the IPD meta-analysis received relevant institutional or country-specific ethics approval, and participants provided written or oral informed consent.

We searched PubMed (https://www.ncbi.nlm.nih.gov/pubmed) and EMBASE (https://www.embase.com) for all available studies up to December 31, 2017 on the association between vitamin D and incident TB disease. Supplementary table 1 provides details of our search strategy (appendix p 3).

#### Study Eligibility and Inclusion Criteria

We placed no restrictions on language. We included all prospective longitudinal studies of human participants at risk for TB if the study measured vitamin D levels in baseline blood samples obtained prior to a diagnosis of TB disease. Studies were included if they confirmed TB diagnosis by microbiological criteria or specified their clinical criteria for ascertaining TB disease and if they collected data on age and gender. We considered any clinical criteria for TB disease based on physician assessment, imaging studies or national guidelines and any laboratory method for assessing vitamin D levels. We did not require studies to control for possible confounders. We also considered reports from conference abstracts. We excluded the following: case reports; animal or in vitro studies; case-control or cross-sectional studies that measured vitamin D levels after TB diagnosis; studies that did not report vitamin D levels; studies of other diseases or non TB-related outcomes; studies of vitamin D and TB treatment outcomes or TB infection or TB immune reconstitution inflammatory syndrome (IRIS); reviews; meta-analyses; letters; editorials; and protocols.

#### Data Collection and Data Items

For each eligible study, two reviewers (OA and SC) independently performed full text reviews and extracted the following information: first author’s last name, year of publication, study design and study aim, country of study, calendar years of study, length of follow up, number of incident TB cases, total number of subjects analyzed, criteria for diagnosing TB disease, laboratory method of vitamin D assay, method of categorizing vitamin D, assessment of HIV status, and covariates in multivariate analyses. Discrepancies were resolved by consensus.

We contacted all authors of the eligible studies identified from the systematic review and requested individual participant data. Lead investigators from all studies agreed to provide data, which were deidentified prior to transfer via email. We requested all available data on possible confounders of the association between vitamin D and TB disease including: age, gender, HIV status, weight, height, BMI, isoniazid preventive treatment, baseline tuberculin skin test result, TB disease history, and comorbid diseases. We also requested baseline vitamin D levels, incident TB disease status during study follow up, time from enrollment to TB diagnosis, and index case smear status if applicable. We reviewed data for consistency with published data and contacted authors for clarifications or missing information as needed.

#### Statistical Analysis

We conducted a one-stage IPD meta-analysis combining data from eligible studies and the Lima cohort study. We used the unified criteria described above to define VDD (25–(OH)D < 50 nmol/L) and insufficiency (25–(OH)D 50–75 nmol/L). We further defined “very low” vitamin D levels as less than 25nmol/L. We considered datasets from each single-country study and from each country within a multi-country study as independent data sources. We used generalized linear mixed univariate and multivariate models to evaluate the association between baseline vitamin D deficiency and risk of incident TB, including an indicator for each independent dataset as a random effect to account for within-study correlation. Multivariate models were adjusted for age, gender, BMI, and HIV. Given sparse data on other variables, we did not adjust for other potential confounders. We also separately evaluated the association between very low vitamin D levels, compared to sufficient levels, and TB disease risk. To determine whether the effect of vitamin D on incident TB differed by HIV status, we conducted a stratified analysis. In a sensitivity analysis, we restricted the main analysis to incident TB cases diagnosed at least 60 days after enrollment.

The IPD meta-analysis was conducted using the R package “lme4”.^16^

## Results

### Lima Cohort Study

Among 6751 HIV-negative household contacts with baseline blood samples, 258 developed TB disease, 66 within 15 days of enrollment and 192 thereafter. Among these 192 secondary TB cases, 152 (79.1%) were microbiologically confirmed, and viable blood samples were available for analysis for 180 (93.8%) at the end of follow-up. Table 1 lists baseline characteristics of the incident cases and their matched controls. The median levels of vitamin D at baseline were similar among cases (53·9 nmol/L; IQR 42·7 – 64·0 nmol/L) and controls (54·7 nmol/L; IQR 44·5 – 67·1 nmol/L; p 0·32) [Table 2]. Median vitamin D level during spring/summer (57·2 nmol/L; IQR 46· 1–69· 0 nmol/L) was higher than during winter (49· 6 nmol/L; IQR 40·5–59·9 nmol/L; p < 0·001).

**Table 1.** Baseline characteristics of participants in Lima cohort study.

|  | **Cases**<br>**(N = 180)**<br>**n (%)** | **N\***<br>**(Cases)** | **Controls**<br>**(N = 709)**<br>**n (%)** | **N\* (Controls)** | **Non-Matched**<br>**household contacts**<br>**(N = 5796)**<br>**n (%)** | **N\***<br>**(Household**<br>**contacts)** |
| --- | --- | --- | --- | --- | --- | --- |
| Age Categories in years |  |  |  |  |  |  |
| < 10 | 4 (2.2) |  | 16 (2.3) |  | 244 (4.2) |  |
| 10 to 19 | 50 (27.8) |  | 200 (28.2) |  | 1024 (17.7) |  |
| ≥ 20 | 126 (70.0) |  | 493 (69.5) |  | 4528 (78.1) |  |
| Male | 94 (52.2) |  | 366 (51.6) |  | 2413 (41.6) |  |
| BMI Categories<sup>†</sup> |  | 179 |  | 706 |  | 5745 |
| Underweight | 8 (4.5) |  | 6 (0.9) |  | 64 (1.1) |  |
| Overweight | 45 (25.1) |  | 299 (42.4) |  | 2881 (50.2) |  |
| Normal | 126 (70.4) |  | 401 (56.8) |  | 2800 (48.7) |  |
| Socioeconomic Status<sup>‡</sup> |  | 171 |  | 698 |  | 5607 |
| Lowest Tertile | 77 (45.0) |  | 228 (32.7) |  | 1847 (32.9) |  |
| Middle Tertile | 66 (38.6) |  | 326 (46.7) |  | 2565 (45.8) |  |
| Highest Tertile | 28 (16.4) |  | 144 (20.6) |  | 1195 (21.3) |  |
| Heavy Alcohol Use<sup>§</sup> | 14 (8.1) | 174 | 64 (9.3) | 692 | 444 (7.9) | 5632 |
| Current Smoking | 13 (7.4) | 176 | 78 (11.2) | 699 | 488 (8.5) | 5711 |
| Self-Reported Diabetes | 6 (3.4) | 179 | 11 (1.6) | 701 | 135 (2.4) | 5739 |
| Comorbid Disease<sup>¶</sup> | 37 (20.6) |  | 175 (24.7) |  | 1380 (23.8) | 5795 |
| Isoniazid Preventive Therapy | 7 (3.9) |  | 108 (15.2) |  | 713 (12.3) | 5790 |
| BCG scar | 159 (88.3) |  | 628 (88.6) |  | 5133 (88.6) | 5795 |
| History of TB | 34 (18.9) |  | 55 (7.8) | 708 | 545 (9.4) | 5782 |
| TB Infection at baseline | 145 (82.4) | 176 | 281 (41.1) | 683 | 2768 (49.0) | 5649 |
| **Index Patient Characteristics** |  |  |  |  |  |  |
| Smear Positive | 156 (86.7) |  | 486 (68.7) | 707 | 4150 (71.6) | 5793 |
| Cavitary Disease | 54 (30.2) | 179 | 175 (25.1) | 697 | 1424 (24.9) | 5722 |

Abbreviation: BCG, Bacillus Calmette–Guérin; BMI, body mass index
\* Total number of subjects with data for corresponding variable.
† We classified adults ≥ 20 years as underweight (BMI < 18.5 kg/m<sup>2</sup>), normal weight (BMI 18.5–< 25 kg/m<sup>2</sup>), and overweight (BMI ≥ 25 kg/m<sup>2</sup>). For children and adolescents < 20 years, we used World Health Organization (WHO) age and gender-specific BMI z-scores tables to classify those with BMI z-score < −2 as underweight and those with z-score > 2 as overweight.
‡ We used principal components analysis of housing asset (housing type, number of rooms, water supply, sanitation facilities, lighting, composition of exterior walls and floor and roof materials) weighted by household size to compute a socioeconomic status score.
§ Self-reported consumption of ≥ 40g or ≥ 3 alcoholic drinks daily.
¶ Heart disease, high blood pressure, asthma, kidney disease, use of steroids or chemotherapy or immunosuppressant, any other self-reported chronic illness

**Table 2.** Baseline levels of vitamin D among cases and controls in Lima cohort study.

|  | **Cases (N = 180)**<br>**median (IQR) or n**<br>**(%)** | **Control (N = 709)**<br>**median (IQR) or n (%)** | **p value \*** |
| --- | --- | --- | --- |
| 25–OH Vitamin D (nmol/L) | 53.9 (42.7 – 64.0) | 54.7 (44.5 – 67.1) | 0.32 |
| Vitamin D Deficient (< 50 nmol/L) | 76 (42.2) | 259 (36.5) | 0.13 |
| Vitamin D Insufficient (50 – 75 nmol/L) | 84 (46.7) | 348 (49.1) | 0.45 |
| Vitamin D Sufficient (> 75 nmol/L) | 20 (11.1) | 102 (14.4) | 1.00 |

\* Univariate p values adjusted for matching factors (age and sex)

In the univariate analysis, household contacts with baseline vitamin D deficiency had a 54% increased risk of incident TB disease compared to those with sufficient levels though this was not statistically significant (95% CI 0·88 – 2·71; p 0· 13) [Table 3]. Vitamin D insufficiency was associated with a smaller increase in risk of TB disease that was not significant (OR 1·23; 95% CI 0·72 – 2·08; p 0·45). After adjusting for BMI categories, socioeconomic status, heavy alcohol consumption, tobacco use, isoniazid preventive therapy, TB history, comorbid disease, self-reported DM, index patient smear status, and season of sample collection, we found baseline vitamin D deficiency and insufficiency remained associated with an increased risk of incident TB disease that was not statistically significant (aOR 1 – 70; 95% CI 0·84 – 3·46; p 0· 14 and aOR 1·25; 95% CI 0·66 – 2.40; p 0·50 respectively) [Table 3]. When we further adjusted for vitamin A deficiency, we continued to find a non-significant increase in risk of TB disease among HHCs with vitamin D deficiency and sufficiency (aOR 1·60; 95% CI 0·76 – 3·39; p 0·22 and aOR 1·21; 95% CI 0·62 – 2·39; p 0·58 respectively).

**Table 3.** Association between vitamin D levels and risk of TB disease among household contacts of TB patients in Lima cohort study.

|  | **Cases/Controls** | **Univariate OR**<br>**(95% CI)**<br>**N = 889** | **p**<br>**value** | **Multivariate OR\***<br>**(95% CI)**<br>**N = 822** | **p**<br>**value** |
| --- | --- | --- | --- | --- | --- |
| Vitamin D Deficient (< 50 nmol/L) | 76/259 | 1.54 (0.88 – 2.71) | 0.13 | 1.70 (0.84 – 3.46) | 0.14 |
| Vitamin D Insufficient (50 – 75 nmol/L) | 84/348 | 1.23 (0.72 – 2.08) | 0.45 | 1.25 (0.66 – 2.40) | 0.50 |
| Vitamin D Sufficient (> 75 nmol/L) | 20/102 | 1.00 |  | 1.00 |  |

\* Adjusted for matching factors (age and sex), body mass index (BMI) categories, socioeconomic status, heavy alcohol consumption, tobacco use, isoniazid preventive therapy, TB history, comorbid disease, self-reported DM, index patient smear status, and season of sample collection.

Our conclusions did not differ from results of the main analysis when we restricted our analyses to cases (and matched controls) diagnosed at least 60 days after index patient enrollment or to microbiologically confirmed TB cases (Table 4).

**Table 4.**
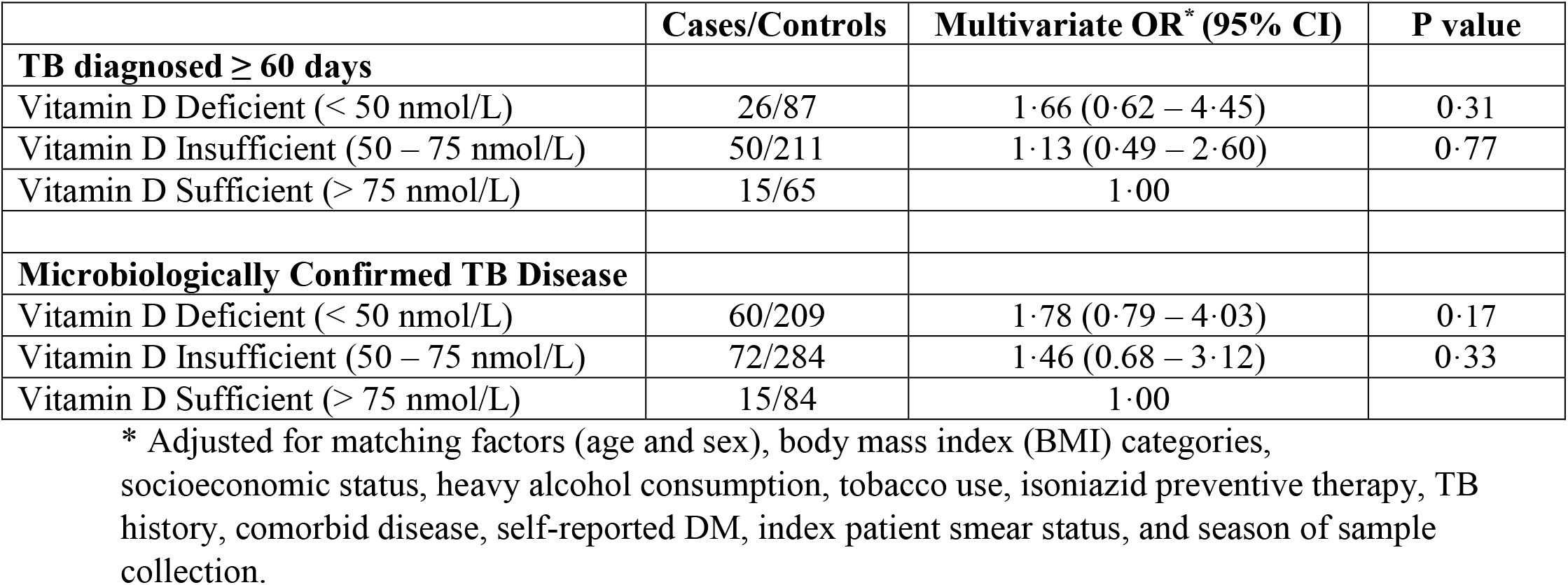
Vitamin D levels and risk of TB disease for TB diagnosed at least 60 days after index case enrollment and microbiologically confirmed TB in Lima cohort study.

### Systematic Review and IPD Meta-analysis

We identified 2689 citations from the PubMed and EMBASE searches. After screening titles and abstracts, we excluded 2678 articles because they were reviews, meta-analyses, letters, editorials or protocols (n = 1212), case reports (n = 515), studies of other diseases or other outcomes (n = 331), animal or *in vitro* studies (n = 247), case-control or cross-sectional studies that assessed vitamin D levels after TB disease diagnosis (n = 159), studies that did not measure vitamin D levels (n = 144), studies of TB treatment outcomes (n = 63), and studies of TB infection (n = 7) [Figure 1]. We reviewed full texts of the remaining 11 articles^17–27^ and further excluded three studies that assessed outcomes of TB-IRIS^24–26^ and one study of TB infection with seasonality of TB.^27^ Table 5 provides information about the seven eligible published studies^17–23^ identified from the systematic review.

**Figure 1.**
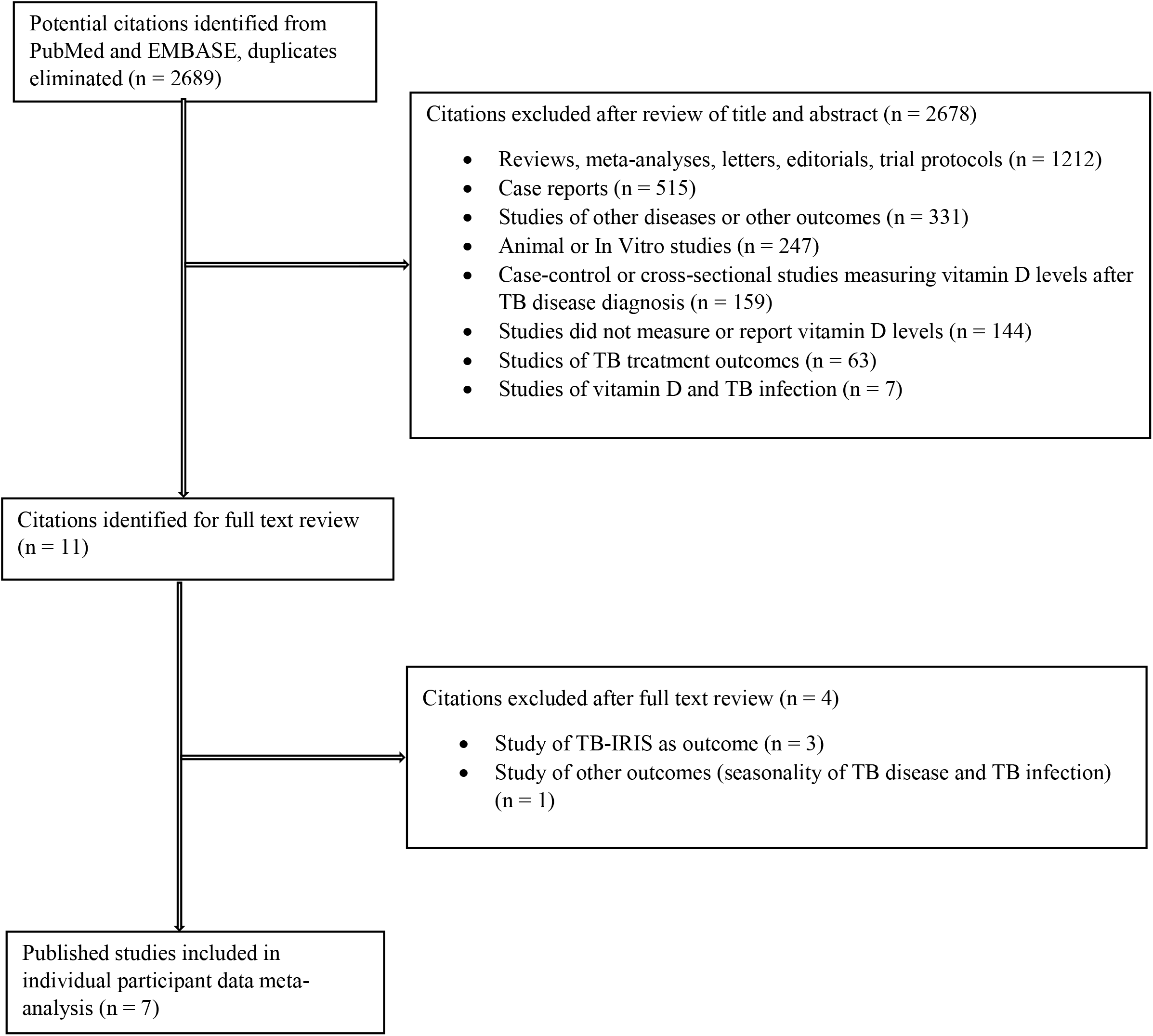
Flow diagram for selection of studies for the individual participant data (IPD) metaanalysis

**Table 5.** Summary of published studies included in the individual participant data (IPD) meta-analysis.

| Study (reference) | Country | Study design and study population | Method of measuring vitamin D | Length of follow up | TB cases/ Total Number of participants | TB disease definition | Adjusted effect estimate (95% CI) |
| --- | --- | --- | --- | --- | --- | --- | --- |
| Arnedo-Pena et al., 2015 (17) | Spain | Prospective cohort study of household and community contacts of TB cases | Electro-chemiluminescence (ECLIA) and Chemiluminescence immunoassays (CLIA) | Mean 1·6 years (± 0·9 years) | 3/523 | Smear or culture positive | aHR for continuous vitamin D: 0·88 (0·80 – 0·97) |
| Gupta et al., 2016 (18) | South Africa | Prospective case-cohort study of HIV positive and HIV exposed infants | Immunoassay | 192 weeks | 100/366 | 2004 South African NTP criteria for definite, probable or possible TB (28) | aHR for vitamin D < 80nmol/L: 1·76 (1·01 – 3·05) |
| Mave et al., 2015 (19) | India | Nested case-control study of HIV positive breastfeeding mothers | Radioimmunoassay | 12 months | 33/120 | Culture confirmed OR Probable TB: 1) smear positive; and 2) histological and clinical features suggestive of TB and response to anti-TB treatment | aOR for vitamin D < 50 nmol/L: 1·57 (0·49 – 4·98) |
| Owolabi et al., 2016 (20) | The Gambia | Prospective cohort study of household contacts of TB cases | ELISA | 24 months | 12/139 | smear/culture positive | Adjusted linear regression estimate for continuous vitamin D: 3·65 (0·59 – 6·71) |
| Sudfeld et al., 2013 (21) | Tanzania | Prospective cohort study of HIV positive patients initiating anti-retroviral therapy (ART) | High Performance Liquid Chromatography | Median 20·6 months (IQR 8·4 – 33·8 months) | 50/1092 | smear positive or CXR | aHR for vitamin D < 50 nmol: 2·89 (1·31 – 7·41) |
| Talat et al., 2010 (22) | Pakistan | Prospective cohort study of household contacts of TB cases | ELISA | 4 years | 8/109 | smear positive or CXR | aHR for 1-log decrement in continuous vitamin D: 5·1 (1·2 – 21·3) |
| Tenforde et al., 2017 (23) | Brazil, Haiti, India, Malawi, Peru, South Africa, Thailand, US, Zimbabwe | Prospective case-cohort study of HIV positive patients initiating ART | Immunoassay | 96 weeks | 70/306 | AIDS Clinical Trial Group (ACTG) Criteria for confirmed, probable or clinical TB (29, Appendix 60) | aHR for vitamin D < 50 nmol: 3·66 (1·16 – 11·51) |

We obtained individual patient data from all eligible studies. One study provided patient data from a multi-site evaluation conducted in nine countries.^23^ Six of the seven studies were prospective cohort or case-cohort studies^17,18,20–23^ while one study was a nested case-control study.^19^ The final combined dataset with our Lima cohort study included 3544 participants from 13 countries: Brazil, The Gambia, Haiti, India, Malawi, Pakistan, Peru, South Africa, Spain, Tanzania, Thailand, US, and Zimbabwe. We analyzed a total of 456 incident TB cases. The median time to TB diagnosis from enrollment was 152·0 days (IQR 44·0 –342·0 days). Table 6 lists the baseline characteristics of all patients analyzed. The majority of the participants (82· 1%) were adults ≥ 18 years of age. HIV status was unknown for 629 (17·7%) patients while 1711 (48·3%) were HIV positive. The median baseline level of 25–OH Vitamin D was 65 0nmol/L (IQR 48·8 – 83·5 nmol/L). The prevalence of vitamin D deficiency at baseline was 26·2% and of “very low” vitamin D levels was 4 8%. Most of the participants with very low vitamin D levels were from studies conducted in India (17·5%),^19^ Spain (25·2%)^17^ and Pakistan (35·7%).^22^

**Table 6.**
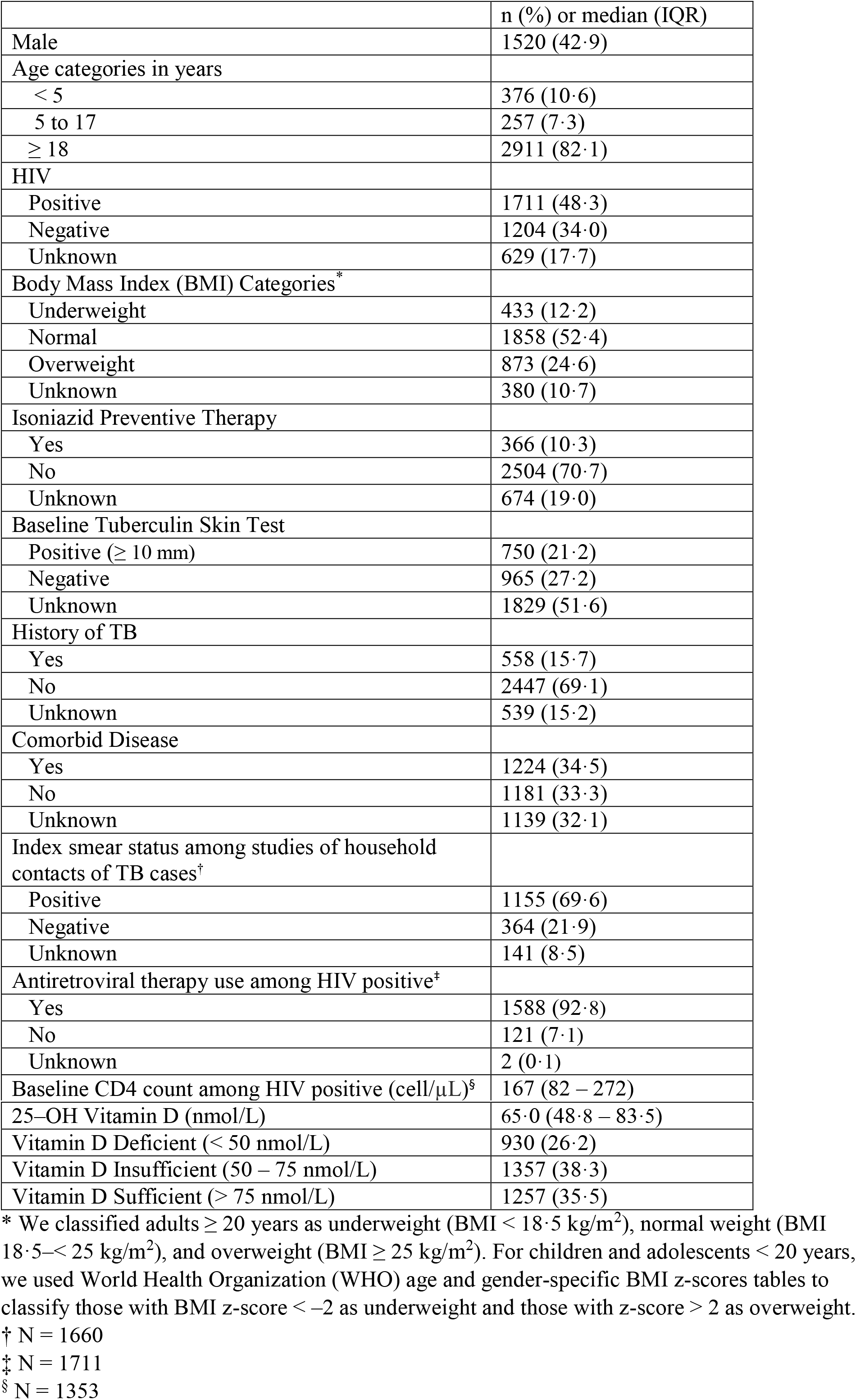
Baseline demographic and clinical characteristics of participants in the individual-participant data (IPD) meta-analysis (N = 3544).

|  | n (%) or median (IQR) |
| --- | --- |
| Male | 1520 (42.9) |
| Age categories in years |  |
| < 5 | 376 (10.6) |
| 5 to 17 | 257 (7.3) |
| ≥ 18 | 2911 (82.1) |
| HIV |  |
| Positive | 1711 (48.3) |
| Negative | 1204 (34.0) |
| Unknown | 629 (17.7) |
| Body Mass Index (BMI) Categories* |  |
| Underweight | 433 (12.2) |
| Normal | 1858 (52.4) |
| Overweight | 873 (24.6) |
| Unknown | 380 (10.7) |
| Isoniazid Preventive Therapy |  |
| Yes | 366 (10.3) |
| No | 2504 (70.7) |
| Unknown | 674 (19.0) |
| Baseline Tuberculin Skin Test |  |
| Positive (≥ 10 mm) | 750 (21.2) |
| Negative | 965 (27.2) |
| Unknown | 1829 (51.6) |
| History of TB |  |
| Yes | 558 (15.7) |
| No | 2447 (69.1) |
| Unknown | 539 (15.2) |
| Comorbid Disease |  |
| Yes | 1224 (34.5) |
| No | 1181 (33.3) |
| Unknown | 1139 (32.1) |
| Index smear status among studies of household contacts of TB cases† |  |
| Positive | 1155 (69.6) |
| Negative | 364 (21.9) |
| Unknown | 141 (8.5) |
| Antiretroviral therapy use among HIV positive‡ |  |
| Yes | 1588 (92.8) |
| No | 121 (7.1) |
| Unknown | 2 (0.1) |
| Baseline CD4 count among HIV positive (cell/μL)§ | 167 (82 – 272) |
| 25–OH Vitamin D (nmol/L) | 65.0 (48.8 – 83.5) |
| Vitamin D Deficient (< 50 nmol/L) | 930 (26.2) |
| Vitamin D Insufficient (50 – 75 nmol/L) | 1357 (38.3) |
| Vitamin D Sufficient (> 75 nmol/L) | 1257 (35.5) |

\* We classified adults $\geq 20$ years as underweight (BMI < 18.5 kg/m<sup>2</sup>), normal weight (BMI 18.5–< 25 kg/m<sup>2</sup>), and overweight (BMI $\geq 25$ kg/m<sup>2</sup>). For children and adolescents < 20 years, we used World Health Organization (WHO) age and gender-specific BMI z-scores tables to classify those with BMI z-score < –2 as underweight and those with z-score > 2 as overweight.
† N = 1660
‡ N = 1711
§ N = 1353

In the univariate analysis, baseline vitamin D deficiency was associated with a 49% increased risk of progression to TB disease (OR 1·49; 95% CI 1·07 – 2·07; p 0·02) [Table 7]. Vitamin D insufficiency also increased the risk of incident TB in a dose-dependent manner (OR 126; 95% CI 0 95 – 1 66; p trend 0 02). Both vitamin D deficiency and insufficiency remained associated with an increased risk of TB disease after we adjusted for age, gender, BMI and HIV status (aOR for deficiency: 1·48; 95% CI 1·04 – 2· 10; p 0·03 and aOR for insufficiency: 1·33; 95% CI 1·00 – 1.77; p 0·05).

**Table 7:** Association between selected baseline characteristics and risk of incident TB disease in the individual participant data (IPD) meta-analysis.

|  | **Univariate OR\***<br>**(95% CI)**<br>**N = 3544** | **P value** | **Multivariate OR†**<br>**(95% CI)**<br>**N = 2769** | **P value** |
| --- | --- | --- | --- | --- |
| Body mass index (BMI) Categories‡ |  |  |  |  |
| Underweight | 1.44 (1.02 – 2.04) | 0.04 | 1.37 (0.96 – 1.96) | 0.08 |
| Normal | Ref |  | Ref |  |
| Overweight | 0.43 (0.32 – 0.58) | <0.001 | 0.43 (0.32 – 0.59) | <0.001 |
| HIV positive§ | 1.44 (0.92 – 2.25) | 0.11 | 1.23 (0.77 – 1.97) | 0.38 |
| Vitamin D Deficient (< 50 nmol/L) | 1.49 (1.07 – 2.07) | 0.02 | 1.48 (1.04 – 2.10) | 0.03 |
| Vitamin D Insufficient (50 – 75 nmol/L) | 1.26 (0.95 – 1.66) | 0.11 | 1.33 (1.00 – 1.77) | 0.05 |
| Vitamin D Sufficient (> 75 nmol/L) | 1.00 |  | 1.00 |  |
|  |  | P trend = 0.02 |  | P trend = 0.03 |

\* Adjusted for age and gender due to presence of age and gender matched case control study in the combined dataset
† Adjusted for age, gender, BMI categories and HIV status
‡ N = 3164
§ N = 2915

When we stratified by HIV status, we found that HIV-positive individuals with vitamin D deficiency were twice as likely to develop TB disease compared to those with normal levels (aOR 2·18; 95% CI 1·22 – 3·90; p 0·01) while HIV-negative participants were only 20% more likely to develop disease (aOR 120; 95% CI 0·75 – 1·94; p 0·44) [Table 8, p for interaction = 0· 17].

**Table 8.** Vitamin D deficiency and risk of incident TB disease stratified by HIV status in the individual participant data (IPD) meta-analysis.^*^

|  | **HIV Positive** |  | **HIV Negative** |  |
| --- | --- | --- | --- | --- |
|  | **Multivariate OR<sup>†</sup>**<br>**(95% CI)**<br>**N = 1576** | **P value** | **Multivariate OR<sup>†</sup>**<br>**(95% CI)**<br>**N = 1193** | **P value** |
| Vitamin D Deficient (< 50 nmol/L) | 2.18 (1.22 – 3.90) | 0.01 | 1.20 (0.75 – 1.94) | 0.44 |
| Vitamin D Insufficient (50 – 75 nmol/L) | 1.32 (0.90 – 1.94) | 0.16 | 1.19 (0.76 – 1.85) | 0.45 |
| Vitamin D Sufficient (> 75 nmol/L) | 1.00 |  | 1.00 |  |
|  |  | P trend =<br>0.01 |  | P trend =<br>0.50 |
| Body Mass Index (BMI) Categories |  |  |  |  |
| Normal | Ref |  | Ref |  |
| Underweight | 1.02 (0.68 – 1.54) | 0.92 | 3.98 (1.79 – 8.85) | 0.001 |
| Overweight | 0.41 (0.21 – 0.78) | 0.01 | 0.45 (0.32 – 0.63) | <0.001 |

\* Excluded datasets from Arnedo-Pena et al. (ref 17) and Talat et al. (ref 22) due to lack of information on HIV status
<sup>†</sup> Adjusted for age, gender and body mass index (BMI) categories

Compared to those with sufficient levels, participant with very low vitamin D levels were twice as likely to develop TB (aOR 2·08; 95% CI 0·88 – 4·92; p trend 0·02) [Table 9]. Among HIV-positive individuals, very low vitamin D levels conferred a 4· 28-fold increased risk of TB (95% CI 0· 85 – 21 44; p trend 0·01). In contrast, very low vitamin D among HIV-negative individuals was associated with a 58% increase in TB disease risk that was not statistically significant (aOR 1·58; 95% CI 0·57 – 4·43; p trend 0·45) [Table 9, p for interaction 0·17].

**Table 9.** Very low vitamin D levels (< 25 nmol/L) and risk of incident TB disease stratified by HIV status in the individual participant data (IPD) meta-analysis.

|  | **Multivariate OR**<br>**(95% CI)** | **P value** |
| --- | --- | --- |
| **All Participants (N = 2769)\*** |  |  |
| Vitamin D < 25 nmol/L | 2.08 (0.88 – 4.92) | 0.10 |
| Vitamin D 25 – < 50 nmol/L | 1.44 (1.01 – 2.06) | 0.04 |
| Vitamin D Insufficient (50 – 75 nmol/L) | 1.33 (1.00 – 1.77) | 0.05 |
| Vitamin D Sufficient (> 75 nmol/L) | 1.00 |  |
|  |  | P trend = 0.02 |
| **HIV Positive (N = 1576)†** |  |  |
| Vitamin D < 25 nmol/L | 4.28 (0.85 – 21.44) | 0.08 |
| Vitamin D 25 – < 50 nmol/L | 2.06 (1.13 – 3.76) | 0.03 |
| Vitamin D Insufficient (50 – 75 nmol/L) | 1.32 (0.90 – 1.94) | 0.15 |
| Vitamin D Sufficient (> 75 nmol/L) | 1.00 |  |
|  |  | P trend = 0.01 |
| **HIV Negative (N = 1193)†** |  |  |
| Vitamin D < 25 nmol/L | 1.58 (0.57 – 4.43) | 0.38 |
| Vitamin D 25 – < 50 nmol/L | 1.18 (0.73 – 1.91) | 0.50 |
| Vitamin D Insufficient (50 – 75 nmol/L) | 1.19 (0.76 – 1.84) | 0.45 |
| Vitamin D Sufficient (> 75 nmol/L) | 1.00 |  |
|  |  | P trend = 0.45 |

\* Model adjusted for age, gender, body mass index (BMI) categories and HIV status
† Model adjusted for age, gender, body mass index (BMI) categories

When we separately considered incident cases diagnosed at least 60 days after enrollment, vitamin D insufficiency remained associated with increased risk of TB disease (aOR 1·42; 95% CI 1·04 – 1·94; p 0·03) while vitamin D deficiency was no longer associated with incident TB (aOR 1·02; 95% CI 0·67 – 157; p 0·91).

## Discussion

The two analyses presented here provide consistent support for a modest dose-dependent effect of vitamin D on future progression of TB disease across multiple studies conducted in diverse contexts. In the IPD, the association of low vitamin D levels with increased TB disease risk is especially strong among HIV-positive individuals whose risk of developing TB was more than four-fold higher for those with very low levels of vitamin D.

Our findings in these human studies support the role of vitamin D in TB disease that has been inferred from more fundamental research which has shown that vitamin D is an important regulator of innate immunity.^4^ *In vitro* studies have enumerated multiple mechanisms by which VDD may influence pathogenesis of TB infection or disease. Vitamin D is implicated in the activation of cathelicidin-mediated killing of ingested mycobacteria,^5,30^ induction of IFN-γ-mediated activity in macrophages,^6^ induction of reactive oxygen and nitrogen species,^31^ stimulation of phagolysosome fusion in infected macrophages,^32^ and inhibition of matrix metalloproteinases involved in the pathogenesis of cavitary pulmonary TB.^33^

In addition to its impact on immunity, vitamin D status has been linked to human metabolic phenotypes that may be involved in the pathogenesis of TB. *In vitro* studies have demonstrated various ways by which vitamin D promotes insulin sensitivity,^34^ and animal models have shown VDD impairs insulin secretion in pancreatic beta cells.^34,35^ Numerous observational studies have also found an inverse association between vitamin D levels and incident type 2 diabetes mellitus (DM).^34^ Given DM is a well described risk factor for TB disease,^36,37^ vitamin D deficiency may also contribute to increased TB risk through its role in modifying risk of diabetes.

Two other lines of evidence point to a possible association between vitamin D and TB. First, multiple studies have reported an association between specific vitamin D receptor (VDR) polymorphisms and TB risk.^8,38^ Although it is not clear that the functional effect of these polymorphisms recapitulates the impact of low vitamin D levels, several studies show that the impact of VDR variants is stronger in the presence of VDD.^18,38,39^ Secondly, TB incidence varies with season and peaks in summer months when vitamin D levels are highest. Some observers have postulated that low levels of sunshine, and hence vitamin D, in winter contribute to an increase in TB infection followed by a rise in TB disease incidence after a six-month lag.^27,40–42^

We also note previous studies have reported that low vitamin A is a strong predictor of incident TB disease.^23,43^ In a previous analysis of the Lima cohort, we found vitamin A deficiency conferred a 10-fold increase in TB disease risk,^11^ and here we show that adjustment for vitamin A modestly attenuates the impact of vitamin D. Similarly, Tenforde et al. also reported that adjusting for vitamin A levels attenuated the effect of vitamin D on TB disease risk.^23^ This raises the possibility that vitamin D levels correlate with other micronutrients implicated in the pathogenesis of TB and these micronutrients may be potent mediators of increased TB risk.

Although we did not detect a statistically significant interaction between vitamin D levels and HIV status, our findings raise the possibility that the effect of low vitamin D on TB risk may be more pronounced among HIV-positive patients. Studies have shown that among HIV-infected individuals, VDD is associated with deleterious immune activation,^44^ lower CD4 counts,^44,45^ higher viral loads,^44^ and accelerated HIV disease progression.^44,46^ Thus, VDD may exacerbate existing immune dysregulation in HIV infection to further increase TB risk or low vitamin D levels may reflect severity of HIV-related immunosuppression. In vitro studies have also demonstrated vitamin D restricts mycobacterial growth in the presence of HIV infection.^47^ A clinical trial is currently underway to evaluate the efficacy of vitamin D supplementation in preventing incident TB among adults with HIV in Tanzania;^52^ the results may help clarify role of vitamin D in HIV-associated TB disease. We also plan to measure inflammatory markers in the Peru cohort to explore association between VDD and immune dysregulation.

We considered possible explanations for why we did not detect a significant association between VDD and TB risk in the Peru cohort, despite its relatively large size. First, since TB incidence is highest in summer^41,42^ and HHCs were recruited when the index case was diagnosed, they are more likely to have been recruited and assessed when their vitamin D levels were highest. If levels later fell and this fall precipitated TB progression, this would not have been detected. Secondly, VDR variants are heterogeneously distributed in different populations and may modify the effect of vitamin D on TB risk. We did not measure VDR variants in Peru and are therefore unable to assess the prevalence of different VDR genotypes in this cohort. The Peru study is also limited by the relatively short (one year) period of follow-up and the fact that it was only powered to detect a three-fold or greater difference in TB incidence among people with VDD.

Our IPD meta-analysis also has some important limitations. Firstly, many possible confounding covariates were not measured across all studies. Therefore, we were unable to account for all possible factors such as baseline infection status, other micronutrient levels and comorbidities that might be associated with both VDD and TB risk. Secondly, although we only examined prospective studies of incident TB disease, the included studies were all observational, and we cannot exclude the possibility that participants had early, undiagnosed TB at baseline that lowered vitamin D levels. Although we addressed this by conducting a sensitivity analysis excluding incident TB cases diagnosed less than 60 days after enrollment, the smaller number of incident cases diagnosed after 60 days reduced the power to detect a statistically significant association. We also cannot exclude the possibility of publication bias if studies with non-significant findings on the link between vitamin D and incident TB have remained unpublished. We did not construct a funnel plot to assess publication bias because we analyzed fewer than ten studies, which used different methods to categorize vitamin D levels and therefore provided effect estimates are not directly comparable. During our systematic review, we attempted to address this by considering data reported from meeting abstracts and none met our inclusion criteria.

In conclusion, in our meta-analysis of prospective studies, we found that vitamin D deficiency was associated with increased risk of incident TB disease. This finding suggests that vitamin D status is a predictor of incident TB disease; however, randomized control trials are needed to determine whether vitamin D supplementation can reduce the risk of developing TB disease. Future research should also elucidate the mechanisms by which low vitamin D influences TB risk.

## Author Contributions

Lima Cohort Study: MBM and MCB led the study design. LL oversaw data collection and management with JTG, RC, ZZ, CC, and RY. RC managed laboratory efforts. MBM supervised data analysis and interpretation in conjunction with OA, C-CH, and MFF.

IPD Meta-Analysis: MBM conceived the study and supervised data extraction, pooling, analysis, and interpretation with C-CH, OA and SC. SA, AA, JB, RB, GD, WWF, DG, MG, BG, AG, NG, RH, JI, NTI, JJ, AK, VM, NM, GM, FMM, OAO, JP, FR, MR, SAS, CRS, MWT, TOT undertook individual studies and patient enrollment.

OA and MBM wrote the first draft of the manuscript, and all authors contributed to manuscript revision.

## Acknowledgments

We would like to thank study participants from all sites.

MBM acknowledges funding from the National Institute of Health (NIH) and National Institute of Allergy and Infectious Diseases (NIAID) [U19AI076217, U01AI057786, TBRU U19AI111224].

OA acknowledges funding from the National Institute on Drug Abuse (NIDA) [T32DA013911] and National Institute of Mental Health (NIMH) [R25MH083620].

AG, GM and SAS acknowledge funding for the NWCS113 International Maternal Pediatric Adolescent AIDS Clinical Trials Group (IMPAACT) study provided by National Institute of Allergy and Infectious Diseases (NIAID) of the National Institutes of Health (NIH) under Award Numbers UM1AI068632 (IMPAACT LOC), UM1AI068616 (IMPAACT SDMC) and UM1AI106716 (IMPAACT LC), with co-funding from the Eunice Kennedy Shriver National Institute of Child Health and Human Development (NICHD) and the National Institute of Mental Health (NIMH). Support of the sites was provided by NIAID and the NICHD International and Domestic Pediatric and Maternal HIV Clinical Trials Network (NICHD contract number N01-DK-9-001/HHSN267200800001C), NIAID [UM1AI069465 to AG] and NINDS [5R01NS077874 to SAS].

MWT and AG acknowledge the NWCS319 project was supported by Award Number U01AI068636 to the AIDS Clinical Trials Group from the National Institute of Allergy and Infectious Diseases and supported by National Institute of Mental Health (NIMH), National Institute of Dental and Craniofacial Research (NIDCR). The work was also supported by the National Institute of Allergy and Infectious Diseases of the National Institutes of Health [U01AI069497, R01AI080417, UM1AI069465 to AG]. The parent trial A5175 was also supported in part by Boehringer-Ingelheim, Bristol-Myers Squibb, Gilead Sciences, and GlaxoSmithKline. MWT also acknowledges funding from NIH [F32AI140511].

AG, VM, and NG acknowledge research reported in this publication was supported by Gilead Foundation, Ujala Foundation (Newton Square PA, USA), and Wyncote Foundation (Philadelphia, PA, USA).

VM, NG, RB, AK and AG acknowledge funding for the SWEN trial was supported by the NIAID [R01 AI45462], the NIH - Fogarty International Center Program of International Training Grants in Epidemiology Related to AIDS [D43-TW0000], the NIAID Byramjee Jeejeebhoy Medical College HIV Clinical Trials Unit [U01 AI069497], and the NIAID’s Baltimore-Washington-India Clinical Trials Unit [UM1 AI069465].

CRS acknowledges funding from the National Institute of Child Health and Human Development [R01 HD32257].

NTI acknowledges funding from the National Commission on Biotechnology (PCST/NCB-AC3/2003), the Higher Education Commission (HEC#20/796/ R&D/06), the International Research Support Initiative Program of the Higher Education Commission Government of Pakistan, and the Bill and Melinda Gates Foundation.

SC acknowledges funding from the NIH Fogarty International Center [K01TW010829].

