## Appendix for "Vitamin D status and risk of incident tuberculosis disease: a systematic review and individual participant data meta-analysis"

### Lima Cohort Study Methods

#### Study Setting and Population

We enrolled a prospective longitudinal cohort of household contacts (HHCs) of index TB patients in Lima, Peru between September 2009 and August 2012. At the time of diagnosis of an index patient, we screened HHCs for TB disease at baseline and referred those with signs or symptoms of TB for diagnosis and treatment according to Peru's national guidelines.[1] Among HHCs without a prior history of TB infection or disease, we assessed baseline TB infection status with the tuberculin skin test (TST). We used a structured questionnaire to obtain clinical, socio-demographic and environmental information from HHCs. We offered all HHCs HIV testing. We also invited all HHCs to provide a baseline blood sample; 60% of HHCs aged 10 years and older complied.

We visited households and re-evaluated all HHCs for pulmonary and extra-pulmonary TB disease at 2, 6 and 12 months after enrollment. We considered HHCs to have incident secondary TB disease if they were diagnosed at least 15 days after index case enrollment and co-prevalent TB disease if they were diagnosed earlier.

At the completion of follow-up, we identified "cases" from among the HHC cohort; these were HIV-negative HHCs with blood samples who developed incident secondary TB disease within one year of follow-up. For each case, we randomly selected four controls from among HHCs who had provided blood samples and who were not diagnosed with TB disease during the study period, matching on gender and age by year.

#### Laboratory Methods

We stored all blood samples at –80ºC from enrollment until end of follow-up. All samples were handled identically, and laboratory personnel were not aware of the case or control status of specimens. Levels of total 25–(OH)D were measured with a commercial competitive enzyme immunoassay kit (Immunodiagnostic Systems Inc., Fountain Hills, AZ), which is sensitive to 5·0 nmol/L. The interassay coefficient of variation for 25–(OH)D ranged from 4·6% to 8·7%. We also measured retinol levels using the high-performance liquid chromatography (HPLC) method described by El-Sohemy et al.[2]

#### Statistical Analysis

We defined vitamin D deficiency (VDD) as serum 25–(OH)D < 50 nmol/L, insufficiency as 50–75 nmol/L and sufficiency as > 75nmol/L.[3] We defined vitamin A deficiency (VAD) as serum retinol < 200 µg/L.[4] We classified adults ≥ 20 years as underweight (body mass index (BMI) < 18·5 kg/m$^2$), normal weight (BMI 18·5–< 25 kg/m$^2$), and overweight (BMI ≥ 25 kg/m$^2$). For children and adolescent HHCs < 20 years, we used WHO age and gender-specific BMI z-scores tables to classify those with BMI z-score < –2 as underweight and those with z-score > 2 as overweight.[5] We classified HHCs as having TB infection at baseline if they reported history of TB disease or positive TST or had a TST result ≥ 10 mm at enrollment.

We used univariate and multivariate conditional logistic regression models to evaluate the association between baseline vitamin D deficiency and risk of TB disease. Multivariate models included baseline covariates identified a priori as potential confounders.

In sensitivity analyses, we repeated our regressions restricting the analysis to people who developed incident TB that was diagnosed at least 60 days after enrollment of index patient and their matched controls. We also performed a sensitivity analysis in which we included only patients with microbiologically confirmed TB and their matched controls. Data were analyzed using SAS v9·4 (SAS Institute, Cary, NC 2013)

### References


1 Peru Ministerio de Salud. Norma Técnica de Salud para el Control de la Tuberculosis. Dirección General de Salud de las Personas. Estrategia Sanitaria Nacional de Prevención y Control de la Tuberculosis. 2006. ftp://ftp2.minsa.gob.pe/descargas/dgsp/ESN-tuberculosis/normaspublicaciones/NTSTBC.pdf (accessed November 16, 2018)

2 El-Sohemy A, Baylin A, Kabagambe E, Ascherio A, Spiegelman D, Campos H. Individual carotenoid concentrations in adipose tissue and plasma as biomarkers of dietary intake. *Am J Clin Nutr* 2002; **76**: 172–179.

3 Institute of Medicine. Dietary reference intakes for calcium and vitamin D. Washington, DC: National Academies Press (US), 2011.

4 Rice AL, West KP Jr, Black RE. Vitamin A deficiency. Global and regional burden of disease attributable to selected major risk factors. Vol 1. World Health Organization, 2004. http://www.who.int/healthinfo/global_burden_disease/cra/en/ (accessed November 16, 2018)

5 World Health Organization (WHO). Child growth standards. 2011. http://www.who.int/childgrowth/software/en/ (accessed November 16, 2018)

**Supplementary Table 1. Search strategy for studies on the association between vitamin D and incident tuberculosis disease**

| | |
|---|---|
| **MeSH Terms for PubMed database search** | "Vitamin D" OR "Vitamin D deficiency" AND "Tuberculosis" |
| **Text Terms for searches on Pubmed and EMBASE databases** | "Vitamin D" OR "Vitamin D deficiency" OR "hypovitaminosis D" OR "25-hydroxyvitamin D" OR "1,25-dihydroxyvitamin D" OR "Vitamin D2" OR "Vitamin D3" OR "Ergocalciferol" OR "Cholecalciferol" AND "Tuberculosis" |
